# Microglial TNFα orchestrates brain phosphorylation during the sleep period and controls homeostatic sleep

**DOI:** 10.1101/2022.03.24.485623

**Authors:** Maria J Pinto, Léa Cottin, Florent Dingli, Victor Laigle, Luís F. Ribeiro, Antoine Triller, Fiona J Henderson, Damarys Loew, Véronique Fabre, Alain Bessis

**Affiliations:** Institut de Biologie de l’École normale supérieure (IBENS), École normale supérieure, CNRS, INSERM, Université PSL, 75005 Paris, France; Institut Curie, PSL Research University, Centre de Recherche, Laboratoire de Spectrométrie de Masse Protéomique, 26 rue d’Ulm, Paris 75248 Cedex 05, France; Center for Neuroscience and Cell Biology (CNC), Institute for Interdisciplinary Research (IIIUC), University of Coimbra, Coimbra, Portugal; Neurosciences Paris Seine - Institut de Biologie Paris Seine (NPS - IBPS), CNRS, INSERM, Sorbonne Universités, Paris, France

## Abstract

The time we spend asleep is adjusted to previous time spent awake, and therefore believed to be under tight homeostatic control. Here, we establish microglia as a new cellular component of the sleep homeostat circuit. By using quantitative phosphoproteomics we demonstrate that microglia-derived TNFα controls thousands of phosphorylation sites during the sleep period. Substrates of microglial TNFα comprise sleep-promoting kinases and numerous synaptic proteins, including a subset whose phosphorylation status codes sleep need and determines sleep duration. As a result, lack of microglial TNFα attenuates the build-up of sleep need, as measured by slow wave activity, and prevents immediate compensation for loss of sleep. Together, we propose that microglia control sleep homeostasis by releasing TNFα that acts at the neuronal circuitry through dynamic control of phosphorylation.

## Introduction

The neuronal circuitries and the neuromodulators regulating sleep are well characterized (*1*, *2*). On the contrary, the molecular mechanisms that control sleep have only been recently investigated. In particular, protein phosphorylation has been described as a fundamental molecular process of sleep control (*3*). The central involvement of phosphorylation in sleep regulation is supported by the fact that kinases, such as SIKs, CaMKIIα/β and ERK1/2 have significant sleep-inducing effects (*4*–*7*). Accordingly, hundreds of proteins, including these kinases, were found to undergo sleep-dependent cycles of phosphorylation/dephosphorylation (*8*, *9*). Because phosphorylation is a major form of regulation of neuronal functions, these changes in phosphorylation were proposed to adjust neuronal functions to sleep or wake brain states. In particular, the dynamics of phosphorylation was shown to principally affect synaptic proteins and to be coupled to synaptic activities (*9*). Phosphorylation of synaptic proteins has also been linked to sleep homeostasis (*10*), the physiological process that guarantees a sufficient amount of sleep. Homeostatic control of sleep is believed to rely on sleep-promoting substances, factors that accumulate in the brain proportionally to the length of the previous wake period and then induce sleep (*11*, *12*). Candidate factors for the homeostatic control of sleep, including adenosine, prostaglandin, TNFα and interleukin-1 (*11*–*14*), have been identified for many years but their molecular mechanisms of action remain poorly understood.

TNFα is a signaling molecule known to control activation of phosphorylation pathways (*15*–*17*). In the brain, TNFα is mostly if not exclusively produced by microglia (*18*, *19*), which are active sensors of brain state (*20*–*22*) and have been recently proposed to participate in the regulation of sleep (*23*–*25*). We therefore hypothesized that microglia-derived TNFα is involved in phosphorylation-based control of sleep. In this study, we demonstrate that microglial TNFα massively modulates cortical phosphorylation exquisitely during the sleep period. We further show that phospho-substrates of microglial TNFα comprise sleep-promoting kinases and numerous synaptic proteins, including those controlling sleep need (*10*, *26*). Consistently, mice with microglial TNFα deletion display altered sleep homeostasis. Our findings reveal microglia as a new cellular actor in the homeostatic control of sleep and offer the first insights into the molecular processes downstream TNFα in sleep modulation.

## Results

### Daily changes in phosphorylations are microglial TNFα-dependent

TNFα has long been identified as a sleep factor (*27*–*29*). In the brain parenchyma, TNFα expression is restricted to microglia (*18*, *19*) and peaks at the sleep period (*30*, *31*), leading us to anticipate a fundamental role of microglia-derived TNFα in the control of sleep and/or sleep-related neuronal network adaptations.

In the brain, the sleep – wake cycle is associated with daily oscillations of phosphorylation (8, 9). To assess the involvement of microglial TNFα in daily changes of protein phosphorylation, we performed quantitative phosphoproteomic analysis of frontal cortex lysates at ZT6 (middle of the sleep period: S) and ZT18 (middle of the wake period: W; fig. 1A) using phosphopeptide enrichment coupled with label-free quantitative mass spectrometry in transgenic control (CTL) and microglia-specific TNFα deleted mice (micTNFα-KO) (fig 1A and fig S1). We focused on the frontal cortex due to its high predominance of sleep slow wave activity (SWA;*32*, *33*), which reflects the homeostatic component of sleep (*12*, *34*). As expected, the sleep – wake cycle was associated with changes in protein phosphorylation. In control animals, 271 phosphosites differed between S and W, representing 2.3% of the phosphosites identified (fig. 1B and table S1). Strikingly, most of this modulation depends on microglial TNFα since 70.7% of these phosphosites showed no more significant changes between S and W in micTNFα-KO. Moreover, half of the phosphosites that still change showed a different direction of change in CTL and micTNFα-KO (fig. 1D), further supporting a role of microglial TNFα in their regulation.

**Figure 1.**
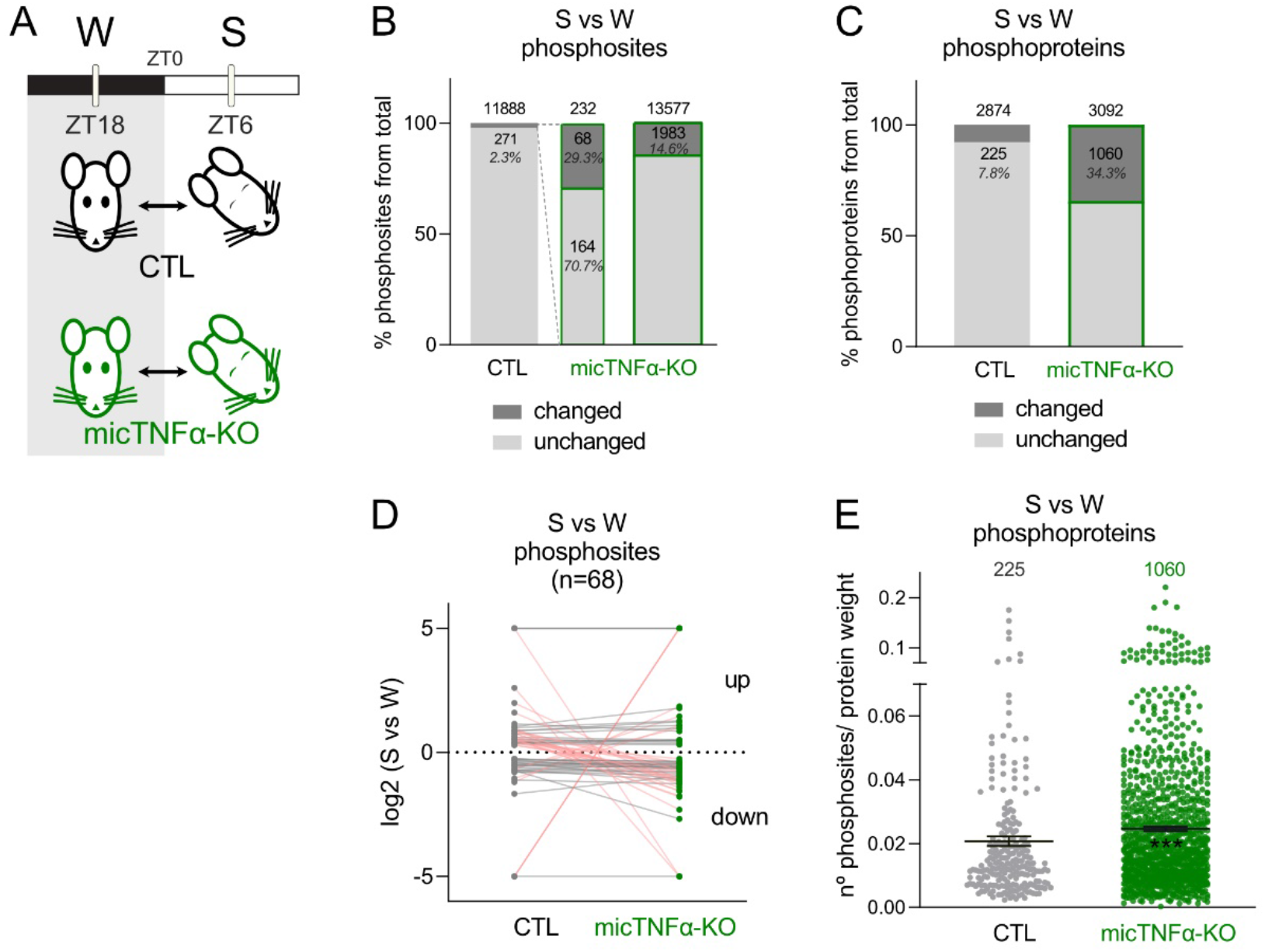
Daily oscillations in cortical phosphoproteome are microglial TNFα-dependent. (**A**) Quantitative analysis of cortical phosphoproteome changes between ZT18 (wake period: W) and ZT6 (sleep period: S) in controls (CTL) and microglia-specific TNFα deleted mice (micTNFα-KO). (**B**) Percentage of phosphosites changing in abundance between S and W in CTL and in micTNFα-KO mice. (**C**) Percentage of phosphoproteins with at least one changing phosphosite in S vs W in CTL and in micTNFα-KO mice. (**D**) S vs W fold change for each phosphosite that changed in both CTL and micTNFα-KO mice. Datapoints above and below the dotted line show up- and down-regulation during S, respectively. Each dot connected by a line represents one phosphosite. Pink lines correspond to phosphosites with a different direction of change in CTL and micTNFα-KO. (**E**) Density of S-W oscillating phosphosites per protein in CTL and micTNFα-KO mice. For each phosphoprotein (represented by a dot), graph shows the number of significantly changed phosphosites in S vs W comparison per protein weight. Average ± SEM is shown. ***p<0.001, Mann Whitney test.

We next compared the phosphoproteome of micTNFα-KO cortex in W and S. Unexpectedly, in contrast to the sparse changes observed in CTL, we found that 14.6% of the identified phosphosites significantly changed between S and W in micTNFα-KO (fig 1B and table S1). Modulation by phosphorylation can occur at multiple sites of the same protein leading us to analyze changes at the protein level. We found that 7.8% of the identified phosphorylated proteins harbor oscillating phosphosites in CTL but as much as 34.3% of the proteins displayed S vs W modulations in micTNFα-KO (fig. 1C). Moreover, phosphoproteins altered by lack of microglial TNFα during the S-W cycle have a higher density of changing phosphorylations than CTL (fig. 1E), suggesting that microglial TNFα massively dampens variations in phosphorylation.

Altogether, these data demonstrate that microglial TNFα is a major regulator of protein phosphorylation during the sleep-wake cycle by acting both as a controller and a buffer of changes in the cortical phosphoproteome.

### Microglial TNFα modulates phosphorylation during the sleep period

We next investigated if the daily control of the phosphoproteome by microglial TNFα occurred preferentially during wake or sleep period by comparing micTNFα-KO and CTL mice phosphoproteomes at W and S (fig 2A). During the wake period, only 2.1% of 11732 phosphosites were regulated by microglial TNFα (fig 2B). In contrast as much as 25.2% of 13731 phosphosites were regulated by microglial TNFα during the sleep period (fig 2B and table S2). Both increases and decreases in phosphosite relative abundance were observed showing that microglial TNFα bidirectionally controls the phosphoproteome (fig 2C). Moreover, during the sleep period, more than 80% of the phosphosites changed more than 2-fold with hundreds of sites changing more than 5-fold (fig 2C and table S2). At the protein phosphorylation level, such TNFα-dependent regulation of phosphorylation affected 7.4% and 48.3% of phosphoproteins during wake and sleep periods, respectively (fig 2D). Of note, at the protein expression level, a similar fraction of the proteome differed between micTNFα-KO and CTL during the wake (7.7%) and the sleep period (6%), with no change above 2-fold (fig S2A and table S3). The reduced overlap between microglial TNFα-dependent proteome and phosphoproteome (fig S3) further suggests that phosphomodulation by TNFα during the sleep period occurs independently from changes at the protein expression level. Our results show that microglial TNFα acts specifically and massively during the sleep period to modulate cortical phosphorylation.

**Figure 2.**
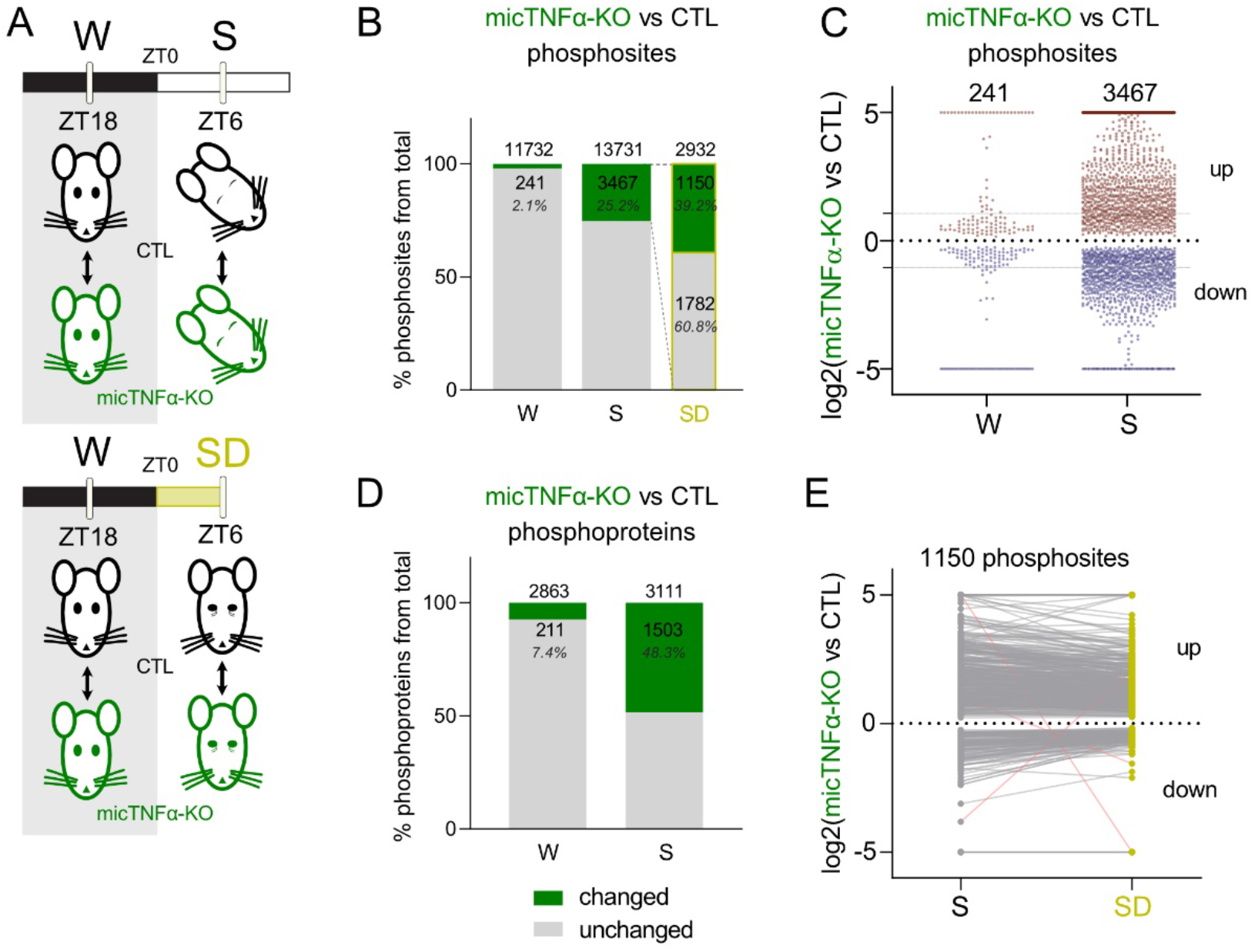
Microglial TNFα acts during the sleep period to modulate the cortical phosphoproteome. (**A**) Quantitative analysis of changes in cortical phosphoproteome between controls (CTL) and microglia-specific TNFα deleted mice (micTNFα-KO) at ZT18 (wake period: W), ZT6 (sleep period: S) and ZT6 after sleep deprivation from ZT0 to ZT6 (SD; yellow). (**B**) Percentage of phosphosites changing significantly in micTNFα-KO vs. CTL comparison during W, S and SD. (**C**) Fold change of significantly different phosphosites in micTNFα-KO vs. CTL comparison at W and S. Each point represents one phosphosite. Datapoints above and below the dotted line show up- and downregulation in micTNFα-KO, respectively. Grey lines indicate 2-fold change. (**D**) Percentage of phosphoproteins with at least one changing phosphosite in micTNFα-KO vs CTL comparison during W and S. (**E**) Comparison of micTNFα-KO vs. CTL fold change for phosphosites changing at both S and SD. Datapoints above and below the dotted line show up- and downregulation in micTNFα-KO, respectively. Each dot connected by a line represents one phosphosite. Pink lines correspond to phosphosites with a different direction of change at S and SD.

Sleep has been shown to trigger daily cycles of phosphorylations (*9*). On the other hand, TNFα expression in microglia is upregulated during the light phase (*31*). We therefore asked whether modulation of the phosphoproteome by microglial TNFα at the sleep period is dependent on sleep mechanisms or is driven by the dark/light cycle. This was assessed by sleep deprivation (SD) experiments. CTL or micTNFα-KO mice were sleep-deprived for 6 hours (from ZT0 to ZT6: SD) and differences in their cortical phosphoproteome were compared to those of mice with *ad libitum* sleep (S) at the same time of day (ZT6; fig 2A). From the pool of microglial TNFα-dependent phosphosites at S (green in fig 2B), 60.8% (1782 out of 2932) did not change between micTNFα-KO and CTL upon SD (fig 2B). This demonstrates that modulation of these phosphosites by microglial TNFα is sleep-dependent. The remaining 39.2% of microglial TNFα-dependent phosphosites (1150 out of 2932 phosphosites) still differed between micTNFα-KO and CTL upon SD (fig 2B) with similar amplitudes and directions of change (fig 2E), demonstrating that modulation of these phosphorylation events by microglial TNFα occurs specifically in the light phase irrespective of sleep. Taken together, these data show that during the sleep period, microglial TNFα modulates the cortical phosphoproteome in a manner dependent on the dark/light cycle and on sleep-intrinsic mechanisms.

### Kinases and phosphatases acting downstream microglial TNFα during the sleep period

The large-scale regulation of the phosphoproteome by microglial TNFα during the sleep period implies an active involvement of kinases and phosphatases. To dissect the molecular mechanisms downstream microglial TNFα, we analyzed the protein levels and phosphorylation of kinases and phosphatases in the proteome of CTL and micTNFα-KO mice during sleep and wake periods. First, we confirmed that the abundance of kinases and phosphatases minimally differed between these mice in W and S (fig S4A). Then, we showed that during the wake period only 3.1% of phosphatases and 6.7% of kinases displayed differential phosphorylation status between micTNFα-KO and CTL mice (fig 3A). In contrast, during the sleep period more than half of phosphatases (56.5%) and kinases (52.5%) underwent TNFα-dependent phosphorylations (fig 3A, S4B). For instance, the phosphatases Plppr4 and Ppm1h have high density of phosphorylation sites regulated by microglial TNFα and so likely prone to its regulation during the sleep period (fig 3B). Furthermore, several regulatory subunits of phosphoprotein phosphatases were identified (e.g. Ppp1r1b, Ppp1r1a; fig 3B, table S2), which possibly account for an indirect modulation of phosphatases’ activity (*35*). Amongst the kinases regulated by microglial TNFα during sleep, we found several known sleep-promoting kinases like CaMKIIα/β (*4*) and MAPKs (*5*, *36*) and the sleep-need associated kinases MARKs (*10*) (fig 3C). We also found that phosphorylation sites known to control the activity of kinases are regulated by microglial TNFα during the sleep period, such as Thr^286^ on CaMKIIα, Tyr^205^/Tyr^185^ on MAPK3/1 and Ser^21^ on GSK3α (table S2) (*37*, *38*). Finally, we found a substantial number of members of the CAMK family (e.g. CaMKIIα/β, Dclk1, Brsk1/2 and MARK1-4) and the CMGC family (including CDKs, MAPKs and GSK3α) displaying multiple sites of microglial TNFα phosphomodulation (fig 3C, table S2). These results emphasize the uncharted potential of microglial TNFα to modulate the activity of kinases and phosphatases through phosphorylation during the sleep period.

**Figure 3.**
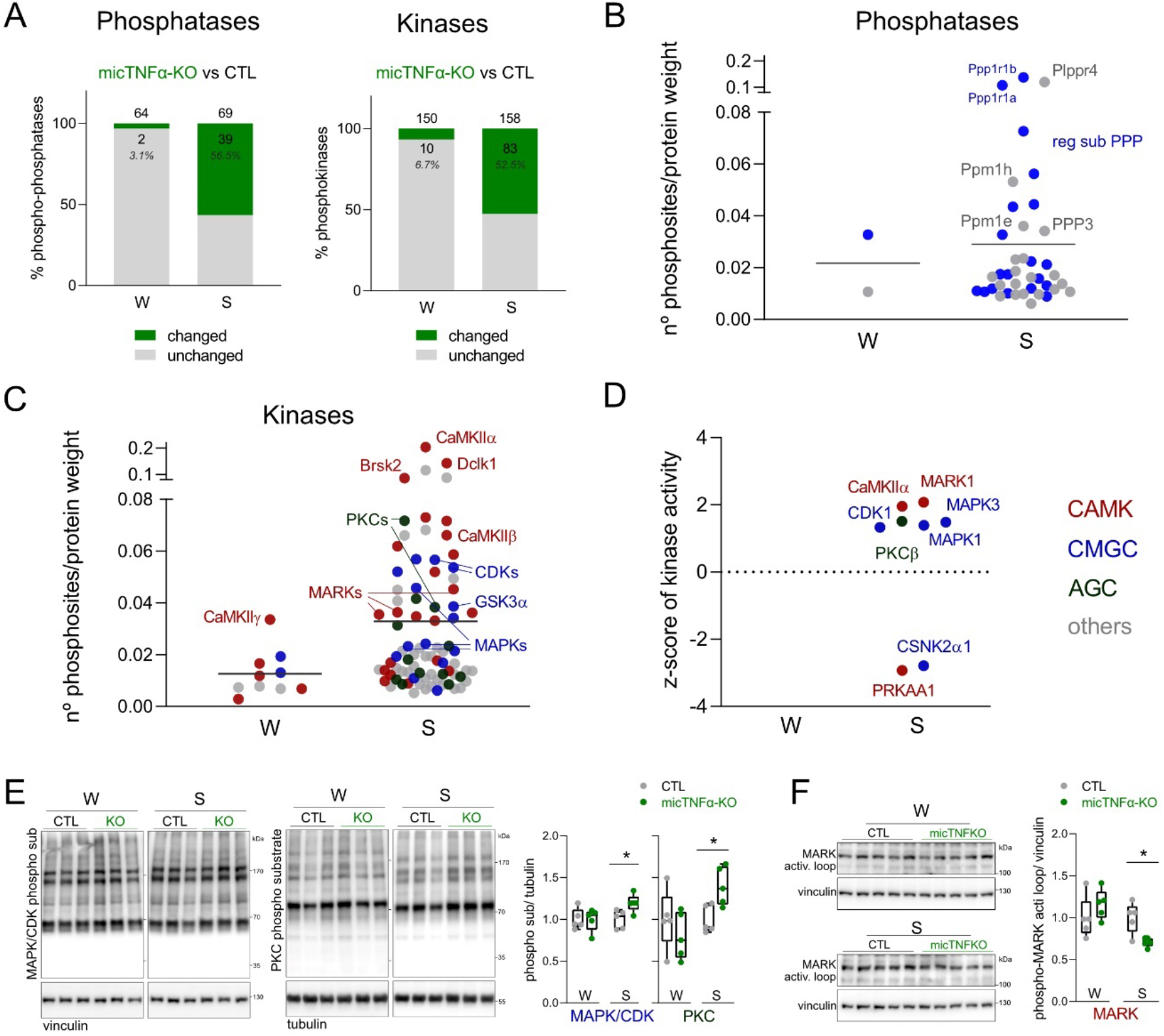
Kinases involved in microglial TNFα phosphomodulation during the sleep period. (**A**) Phosphoregulation of phosphatases and kinases by microglial TNFα. Graphs show percentage of phospho-phosphatases and phosphokinases with at least one changing phosphosite in micTNFα-KO vs CTL comparison at W and S. (**B, C**) Density of phosphosites on (**B**) phosphatases and (**C**) kinases modulated by microglial TNFα at W and S. Graphs show the number of significantly changed phosphosites in micTNFα-KO vs CTL comparison per protein weight. Each dot is one phosphoprotein. Phosphatases are shown in grey and regulatory subunits of phosphoprotein phosphatases (reg sub PPP) in blue. Kinases are color-coded according to the major families: calcium/calmodulin regulated kinases (CAMK, red); CDK, MAPK, GSK3 and CLK kinases families (GMGC, blue); protein kinase A, G and C families (AGC, dark green). (**D**) Kinases with altered activity between controls and micTNF-KOs predicted by robust kinase activity inference (RoKAI). Graph shows the z-score value of each kinase assigned by RoKAI. Kinases are color-coded according to kinase group. (**E, F**) Validation by western blot of altered activity of predicted kinases between CTL and micTNF-KOs. (**E**) *Left*, Immunoblots of cortical lysates using antibodies specific to MAPK/CDK and PKC target phosphorylation motifs. *Right*, Ratio between phosphorylated substrates and loading control normalized to CTL at W and S. (**F**) *Left*, Immunoblots show signal of phosphorylated MARK activation loop. *Right*, Ratio between phosphorylated MARK activation loop and loading control normalized to CTL at W and S. (**E, F**) n=5 mice per group. *p<0.05, Mann Whitney test.

We then used RoKAI, an unbiased kinase activity inference tool to identify kinases with changed activity based on changes in the phosphorylation profile of their substrates (*39*). RoKAI identified mostly members of the CAMK and CMGC family (fig 3D). Notably, MARK1, CaMKIIα, PRKAA1, CDK1, MAPK1/3 and CSNK2α1 were identified as kinases with altered activity during the sleep period in micTNFα-KO as compared to CTL (fig 3D). PKCβ, a member of the AGC family was also identified as a kinase putatively regulated by microglial TNFα during sleep. Of note, no significant hits were obtained at wake time (fig 3D). We then validated the RoKAI predictions by western blot analysis and we chose MAPKs, CDKs, PKCs and MARKs as test cases. As expected, an increase in the phosphorylation of substrates of MAPK/CDK and PKC was observed on micTNFα-KOs frontal cortex proteins in comparison to CTL during the sleep period but not during the wake period (fig 3E). Similarly, reduced phosphorylation of the activation loop of MARKs, known to correlate with kinase activity (*40*), only occurs at sleep time in micTNFα-KO (fig 3F). We further verified that the phosphorylation of substrates of PKA and AKT, whose activities were not predicted to be affected (fig 3D), did not display changes in micTNFα-KO as compare to CTL mice (fig S5). Finally, we used organotypic slices to demonstrate acute effects of TNFα on the candidate kinases. This confirmed that short-term neutralization of TNFα mimicked the changes in MAPK/CDK and PKC signaling pathways observed on micTNFα-KOs *in vivo* (fig S6). Together, these observations support a direct effect of TNFα on the above-mentioned kinases.

### Phosphorylation of sleep-related synaptic proteins is regulated by microglial TNFα

To gain further insight into the roles of microglial TNFα-mediated phosphorylation during the sleep period we used the Reactome Pathway Analysis (*41*, *42*), an unbiased pathways analysis tool, to characterize the functions enriched in the substrates of TNFα phosphomodulation (fig 2C). Strikingly, the enriched pathways were almost exclusively related to synaptic functions both at the presynaptic and postsynaptic level (fig 4A). In agreement, we found that 34.7% (521 out of 1503) of the entire pool of TNFα-regulated phosphoproteins during the sleep period is annotated as synaptic. Moreover, several of these synaptic proteins undergo modulation by microglial TNFα at multiple phosphorylation sites, particularly during the sleep period (fig 4B). These include presynaptic (e.g. synapsins, RIMs, Syngap2, Bassoon, Piccolo) and postsynaptic proteins (e.g. NMDARs, Shanks, Ank2/3, Dlgap1, Agap2), as well as synaptically-localized kinases (e.g. CaMKIIα, Dclk1, MARKs). Finally, no synaptic pathways were enriched when considering the proteins whose abundance is regulated by microglial TNFα during the sleep period (fig S2b). Thus, protein phosphorylation, but not control of protein levels, likely underlies microglial TNFα regulation of synapses during the sleep period.

**Figure 4.**
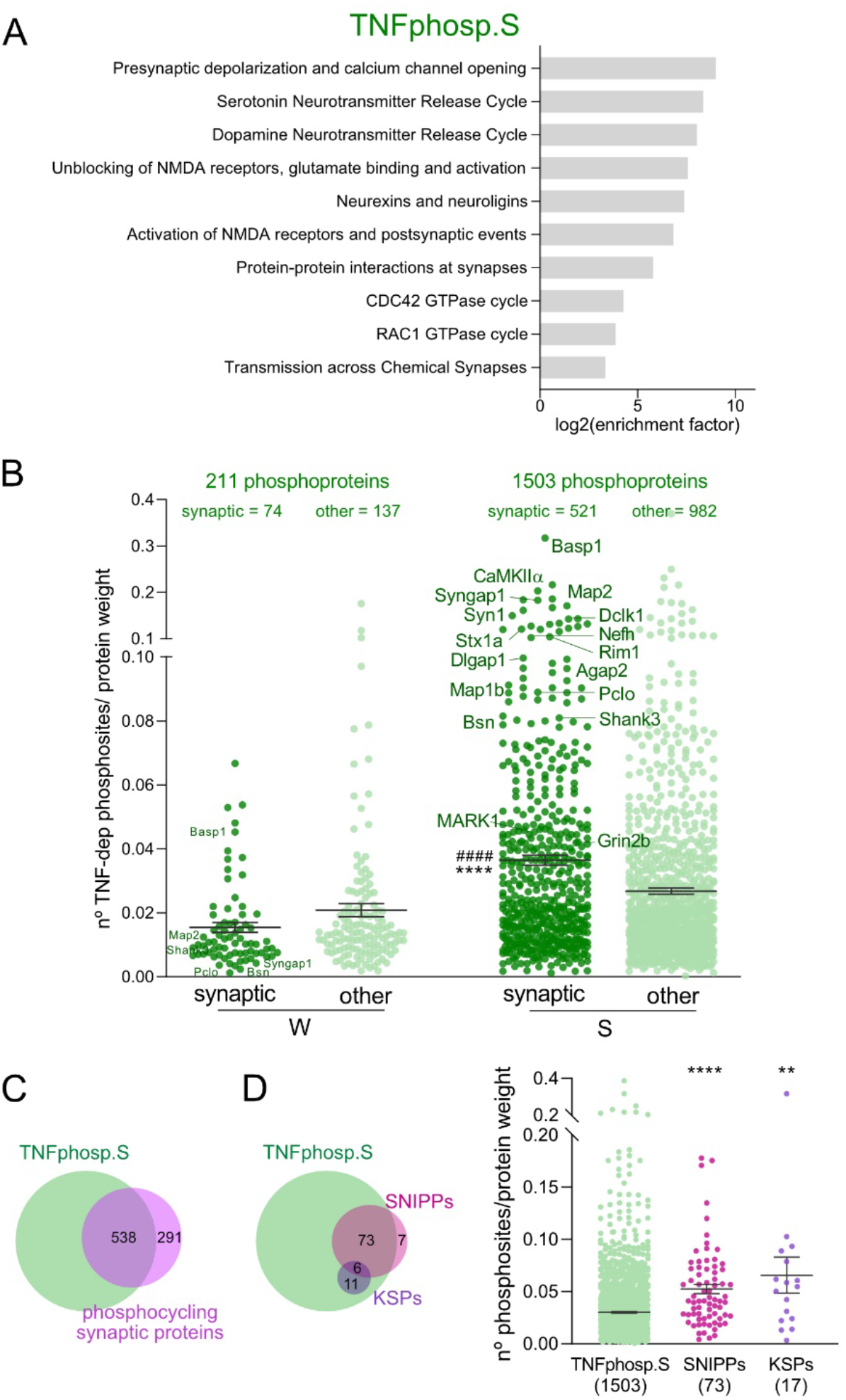
Sleep-associated synaptic proteins are major targets of microglial TNFα phosphomodulation during the sleep period. (**A**) Pathway analysis of substrates modulated by microglial TNFα through phosphorylation during the sleep period (TNFphosp.S). (**B**) Density of phosphosites on synaptic and non-synaptic proteins (other) modulated by microglial TNFα at W and S. Graph shows the number of significantly changed phosphosites in micTNFα-KO vs CTL comparison per protein weight. Each dot corresponds to one phosphoprotein. ****p<0.0001 compared to W synaptic and ^####^p<0.0001 compared to S other, one-way ANOVA. (**C**) Venn diagram shows high overlap of TNFα-modulated phosphoproteins during the sleep period (TNFphosp.S) and synaptic proteins harboring daily phosphorylation changes (*9*). (**D**) *Left*, Venn diagram of microglial TNFα-modulated phosphoproteins during the sleep period (TNFphosp.S) and sleep-associated phosphoproteins SNIPPs (*10*) and KSPs (*26*). *Right*, Density of phosphosites modulated by microglial TNFα at S on all TNFα phosphosubstrates (TNFphosp.S), SNIPPs and KSPs. Graph shows the number of significantly changed phosphosites in micTNFα-KO vs CTL comparison per protein weight. Each dot corresponds to one phosphoprotein. ****p<0.0001 and **p<0.01 compared to TNFphosp.S, one-way ANOVA.

Several studies have demonstrated that phosphorylation of proteins, in particular of synaptic proteins, controls the duration of sleep (*4*, *6*, *10*, *26*). For instance, sleep - wake cycles were shown to specifically drive the phosphorylation of more than 800 synaptic proteins (*9*). Remarkably, we found that 65% of these synaptic proteins are regulated by microglial TNFα in the sleep period (fig 4C). Homeostatic regulation of sleep relies on internal mechanisms that encode sleep need according to the previous time spent awake. The phosphorylation – dephosphorylation of 80 synaptic proteins identified as the synaptic sleep-need index phosphoproteins (SNIPPs) has been proposed to code sleep need (*10*). Phosphorylation of another set of synaptic proteins identified as keystone sleep phosphoproteins (KSPs) has been causally linked to sleep induction (*26*). Remarkably, 91% of SNIPPs and 100% of KSPs are substrates of microglial TNFα phosphomodulation during the sleep period (fig 4D). Furthermore, the high density of phosphosites modulated by microglial TNFα on SNIPPs and KSPs highlights the prime ability of microglial TNFα to regulate their phosphorylation status (fig 4D). Taken together, our results identify microglial TNFα as a major modulator of phosphorylation of sleep-related synaptic proteins and therefore a putative upstream regulator of sleep homeostasis.

### Homeostatic control of sleep requires microglial TNFα

Sleep homeostasis sets the duration and intensity of sleep according to the duration of prior wake. Actually, sleep need increases during spontaneous or enforced wake and then dissipates during the following sleep period. It has been proposed that the phosphorylation of SNIPPs determines sleep need and is therefore part of the regulatory mechanism of sleep homeostasis (*10*). Because we have shown that microglial TNFα massively controls the phosphorylation status of SNIPPs, we hypothesized that microglial TNFα is a regulator of sleep need. This was first shown by measuring dynamic changes in cortical EEG SWA in NREM sleep, a reliable indicator of sleep need (*12*, *34*), in micTNFα-KO and CTL mice throughout a 24-hr sleep-wake cycle (baseline recording) (fig 5A). As expected, in CTL mice SWA increased during the dark (wake) period reflecting the build-up of sleep need, and decreases progressively during the light (sleep) period when sleep need dissipates (fig 5A). Strikingly, at the onset of the light (sleep) period, which is the time of highest accumulation of sleep need, the amount of SWA was significantly lower in micTNFα-KO mice as compared to CTL mice (fig 5A). To further demonstrate a role of microglial TNFα as a regulator of sleep need, sleep homeostasis in micTNFα-KO mice was then challenged by sleep deprivation (SD). Sleep deprivation substantially increases sleep need and triggers a compensatory response by boosting subsequent SWA in NREM sleep and sleep duration (*32*, *43*, *44*). CTL and micTNFα-KO mice were forced to stay awake during their normal sleep period (SD from ZT0 to ZT6) and the consequences of SD were assessed in the subsequent recovery phase. As expected, during the first hours of recovery following SD, CTL mice displayed a sharp increase in SWA in NREM sleep (fig 5A), reflecting elevated sleep need (*12*, *34*). Moreover, SD elicited a NREM sleep rebound at the expense of wake levels during the following recovery phase (fig 5B). This effect was the consequence of an increase in the mean duration of NREM sleep episodes (Table S4). Remarkably, at the beginning of the recovery period micTNFα-KO mice displayed less SWA as compared to CTL mice (fig 5A) consistent with a compromised build-up of sleep need following extended periods of wake. In contrast to CTL mice, micTNFα-KO mice lacked NREM sleep rebound and concomitant decrease in wake amounts during the recovery phase following SD indicative of impaired sleep homeostasis (fig 5B). In addition, REM sleep amounts in micTNFα-KO mice were increased in the Sham condition and decreased following sleep deprivation (fig 5B) suggesting an additional role of microglial micTNFα in REM sleep regulation.

**Figure 5.**
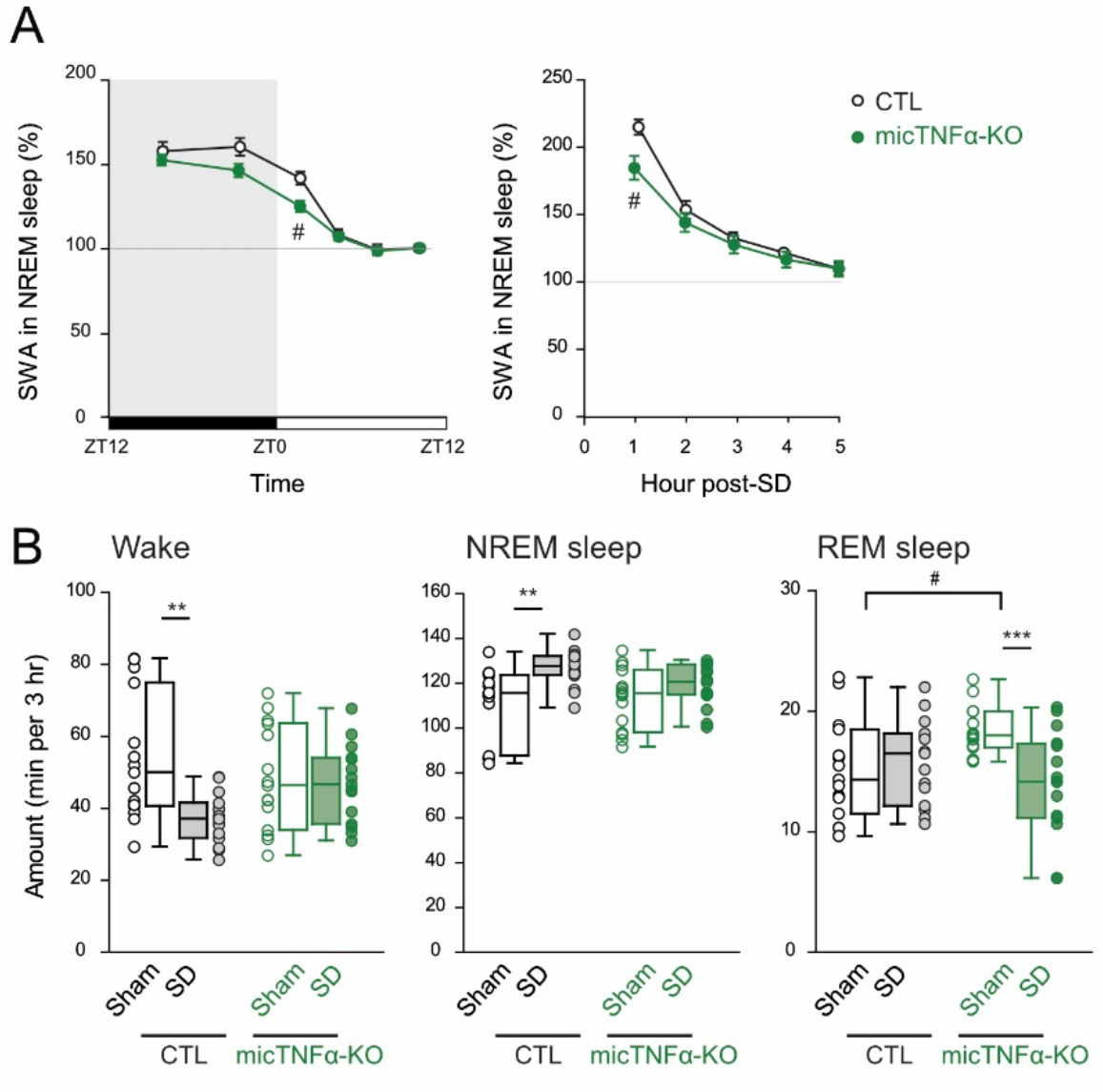
Microglial TNFα modulates sleep homeostasis. (**A**) Time course of SWA during NREM sleep across a 24 hr baseline sleep-wake cycle (left) and in the first 5 hr after sleep deprivation (SD; right) in controls CTL (black) and micTNFα-KO (green) mice. Mean SWA values normalized to the mean SWA during ZT8 to ZT12 of the baseline recording, a period during which sleep need is lowest (see Methods). SWA in baseline sleep: CTL, 14 mice; micTNFα-KO, 15 mice. SWA after sleep deprivation: CTL, 12 mice; micTNFα-KO, 15 mice. #p< 0.05 between CTL and micTNFα-KO, two-way repeated-measure ANOVA followed by Sidak’s multiple comparisons test. (**B**) Wake, NREM sleep and REM sleep amounts in the first 3 hr of the recovery phase after sleep deprivation (SD, plain) or Sham condition (Sham, empty) in CTL (black) and micTNFα-KO (green) mice. SD: mice were sleep deprived for 6 hours starting at ZT0; Sham condition: each mouse was gently awakened at ZT6, 2 days prior to the SD procedure (matched control recording). Data are represented as mean ± SEM. For multiple group comparisons: **p<0.01 and ***p<0.001 Sham versus SD and #p< 0.05 Sham CTL versus Sham micTNFα-KO, two-way repeated-measure ANOVA followed by Sidak’s multiple comparisons test.

Altogether, these results agree with impaired build-up of sleep need in micTNFα-KO mice in response to prolonged wake and confirm that microglial TNFα participate to homeostatic sleep control.

## Discussion

Sleep is under tight homeostatic control but the cellular and molecular mechanisms of such control are still largely unknown. We now demonstrate that microglial TNFα extensively modulates cortical phosphoproteome during the sleep period (fig. 2) including many kinases, phosphatases (fig. 3) and synaptic proteins (fig. 4). A set of these microglial TNFα-targeted substrates are SNIPPs and KSPs (fig. 4), proteins whose phosphorylation status has been proposed to encode sleep need and control sleep duration, respectively (*10*, *26*). In agreement, we showed that microglial TNFα is required for expression of sleep need and concomitantly controls homeostatic sleep rebound (fig. 5).

### Homeostatic regulation of sleep by microglia

The homeostatic regulation of sleep sets the duration and intensity of sleep as a function of the preceding time spent awake. Such control relies on the build-up of sleep need during wake and subsequent translation into sleep drive. One currently held belief is that sleep need arises in discrete neuronal networks driven by the accumulation of sleep-promoting substances following prolonged periods of neuronal activity during extended wake (*11*, *12*). Astrocytes have been implicated in sleep homeostasis via the release of adenosine (*13*, *45*, *46*), which acts on the basal forebrain and hypothalamus to induce sleep (*47*, *48*). Despite work in *Drosophila* suggesting a connection between glia-derived immune factors and homeostatic sleep (*49*, *50*), the involvement of microglia has remained elusive. Our work now provides functional evidence that microglia are active players in sleep homeostasis via TNFα. It is worth noting that homeostatic sleep control by microglia may not be limited to the secretion of TNFα, as these cells control the levels of adenosine (*22*) and can release other known sleep-promoting substances such as interleukin-1 and prostaglandins (*51*, *52*). The somnogenic effect of TNFα has long been described. Central or peripheral injections of TNFα increase NREM sleep (*27*, *29*, *53*), whilst neutralization of TNFα signaling attenuates spontaneous sleep (*28*) and homeostatic sleep rebound in rabbits (*54*). In the brain parenchyma, TNFα expression is restricted to microglia (*18*, *19*, *55*). However, peripheral TNFα can cross the blood brain barrier and play an active role in the neuronal circuitry (*56*, *57*). Assessing the relative contribution of central and peripheral TNFα-producing cells to sleep control is thus an important issue. Herein we prove that, despite not being involved in the control of basal sleep amounts (*58*), TNFα of microglial origin controls NREM sleep recovery when sleep homeostasis is challenged by sleep deprivation (fig. 5).

The involvement of microglia in homeostatic sleep control is consistent with current models of sleep regulation and the well-known ATP-driven microglia behavior. The former posits that accumulation of ATP due to enhanced brain activity during prolonged wake triggers the release of cytokines which in turn promote sleep (*59*, *60*). This concept fits with the ability of microglia to react to neuronal activity via ATP-mediated changes in their behavior (*20*, *22*, *58*). Indeed, ATP triggers the release of TNFα, interleukin-1 and prostaglandin by microglial cells (*61*–*63*). Collectively, our work and these findings support the idea that microglia sense extended periods of wake via ATP and respond by releasing proportional amounts of TNFα, which in turn feedbacks onto the neuronal networks to control sleep. Alternatively, upon extended wake microglia could be recruited by wake-promoting neuromodulators (such as noradrenaline, serotonin, dopamine and histamine), which modulate microglia function including triggering the release of sleep substances like TNFα (*21*, *64*, *65*).

The molecular mechanisms linking sleep-promoting substances and sleep need transformation into sleep drive are poorly known. We now propose that one of the molecular determinants of microglial TNFα on homeostatic sleep regulation is protein phosphorylation in the cortex. Such phosphorylation-mediated control can occur at two different levels: either contributing to the molecular coding of sleep need and/or transforming sleep need into sleep drive. In support of the former possibility, microglial TNFα controls (de)phosphorylation events in SNIPPs (fig. 4), proteins whose phosphorylation status determines sleep need (*10*). For instance, mGluR5 was previously shown to contribute to sleep need (*66*) and our data identifies TNFα-dependent phosphorylation sites that affect mGluR5 signaling (Ser^839^ and Thr^840^ on mGluR5; table S2) (*67*). In agreement, lack of microglial TNFα leads to reduced slow wave activity, a classic measure of sleep need, at times of high sleep pressure (fig. 5). In support of the latter, microglial TNFα also adjusts the phosphorylation of the sleep-inducing phosphoproteins KSPs (*26*) and sleep-promoting kinases like CaMKIIα/β (*4*) and MAPKs (*5*, *36*) (fig. 3 and 4). Phosphosites identified on the mentioned kinases are known to control their activity (table S2), suggesting that microglial TNFα could trigger sleep rebound after sleep deprivation by controlling the activity of specific kinases via phosphorylation. Importantly, despite the hundreds of proteins under microglial TNFα phosphoregulation at the sleep period (table S2), the structure of baseline sleep is not disrupted upon loss of microglial TNFα (*58*). This observation suggests that microglial TNFα-dependent phosphorylation is not required to keep sleep time and duration at steady-state conditions. However, when sleep-wake homeostasis is unbalanced, as in periods of insufficient sleep, control of the phosphoproteome by microglial TNFα is determinant. Finally, given that the phosphoproteomic analysis was performed in the frontal cortex, our work strengthens findings attributing a role for the cortex in sleep homeostasis (*68*–*70*). Microglia, via TNFα-mediated phosphomodulation, likely act in synchrony with discrete cortical circuits to regulate homeostatic sleep.

### A potential role of microglial TNFα in synaptic plasticity during sleep

Protein phosphorylation has a pivotal role in controlling synaptic plasticity (*37*, *71*, *72*), which occurs during sleep to shape neuronal networks (*8*, *73*, *74*). In line with the extensive modulation of cortical phosphoproteome during the sleep period, our data further suggests a promising role of microglial TNFα in orchestrating sleep-associated synaptic plasticity. Indeed, an unbiased pathway analysis to the substrates of microglial TNFα phosphomodulation at the sleep period reveals enrichment of synaptic functions (fig. 4). Notably, many of these substrates change in phosphorylation upon induction of homeostatic downscaling (fig. S7) (*75*), a form of synaptic plasticity believed to occur during sleep to weaken excitatory synapses (*8*, *73*). Furthermore, predictive and experimental validation of microglial TNFα-targeted kinases allowed identification of PKCs and MAPKs (fig. 3), which are known to regulate synaptic plasticity (*76*–*78*).

In further support of a prominent role of microglial TNFα at synapses during sleep, several known functional phosphosites on pre- and postsynaptic proteins were identified as regulated by TNFα during sleep (fig.4 and table S2). For instance, the postsynaptic NMDAR 2A subunit (Grin2a) display changes in the phosphorylations of Ser^929^ and Ser^1459^, residues involved in the trafficking and synaptic presentation of the receptor (*79*, *80*), whilst NMDAR 2B subunit (Grin2b) becomes dephosphorylated at Ser^1303^ known to control its binding to CaMKII (*81*). The latter is a central kinase in postsynaptic plasticity whose activity is controlled by phosphorylation at Thr^286^ (*37*), also shown to be modified by TNFα. Another target phosphosite is Ser^1539^ on the postsynaptic scaffold Shank3, recently shown to control synaptic availability of the protein and thereby expression of homeostatic plasticity (*82*). Examples can also be drawn on regulation of presynaptic proteins by specific phosphorylations, herein identified as microglial TNFα-dependent during sleep (fig.4 and table S2). For instance, release and turnover of synaptic vesicles is controlled by phosphorylation of syntaxins at Ser^14^ and synapsins at Ser^62^ (*83*, *84*) and long-term plasticity is supported by enhanced presynaptic release via Rim1 phosphorylation at Ser^413^ (*85*). Together, our data point toward an active participation of microglial cells to synaptic remodeling during sleep via TNFα-controlled phosphorylation. Supporting findings have recently been reported on sleep-dependent plasticity of inhibitory synapses (*58*).

Finally, we showed that modulation of the phosphoproteome by microglial TNFα occurs specifically during the sleep period downstream a joint action of sleep and time-of-day-dependent processes (fig. 2). Future work is needed to unravel the mechanisms linking microglia sensing of brain states, like sleep, and its translation into molecular processes that tune the neuronal network.

## Materials and Methods

### Animals and housing

Animal experiments were performed in accordance with the European Committee Council Directive 86/609/EEC and procedures approved by the local Charles Darwin Ethical Committee (Ce5-2014-001; 1339-2015073113467359 and 2018022121466547). CX_3_CR1^GFP^ (*86*); CX_3_CR1^CreERT2^ (*87*); TNF^flox^ (*88*) mouse lines were housed at the animal facility of Institut de Biologie de l’ENS or at the animal facility of Institut Biologie Paris Seine (Paris, France). CX_3_CR1^GFP^ and CX_3_CR1^CreERT2^ were kindly provided by Sonia Garel and TNF^flox^ by Etienne Audinat. All mice used were 4- to 5-month-old males, housed under standard conditions (12h light/dark cycle; lights on at 7:00 a.m.) with food and water ad libitum.

### Conditional microglia-specific TNFα deletion

Proteomic and phosphoproteomic experiments were performed on mice following conditional deletion of TNFα on microglia cells. To accomplish so, CX3CR1^GFP/+^:TNF^f/f^ (transgenic controls, CTL) and CX3CR1^CreERT2/+^:TNF^f/f^ (microglia-TNFα depleted mice, micTNF-KO) mice were fed with tamoxifen-containing food (1000mg tamoxifen citrate in 1 kg of chow, formulated by SSNIFF, A115-T71000) for 6 days. Experiments were performed 5 to 6 weeks after tamoxifen feeding, which allows for repopulation of short-lived peripheral CX3CR1^+^ cells thus restricting recombination of the TNFα locus to microglia (*89*). To confirm microglia depletion of TNFα, adult primary microglia were isolated and cultured as previously described (*90*). In brief, PBS-perfused adult mouse brains were dissected and dissociated in 30-40U papain (Worthington, LK003176), 7.2 U dispase II (Sigma, D4693) and 10 mg/ml DNAse (Sigma, DN25). Following mechanical dissociation, microglia were isolated using a Percoll gradient (Sigma, P4937) and plated in DMEM/F12 (GIBCO, 31331-028) supplemented with 10% heat-inactivated fetal bovine serum (GIBCO, 10500-064), 10 U/mL penicillin and 10μg/mL streptomycin. Cells were stimulated with 10μg/mL LPS for 10h to induce TNFα release. Medium was collected and centrifuged for 5 min at 10000g at 4°C. Levels of TNFα in the supernatant were determined by the mouse TNFα uncoated ELISA kit (Invitrogen, 88-7324).

### Tissue collection and sample preparation for mass spectrometry

Following tamoxifen treatment, mice were put in individual cages for 2 weeks before tissue collection. At ZT6 and ZT18, which correspond to the middle of the light and dark phases, mice spend most of their time asleep and awake, respectively. ZT6 and ZT18 timepoints were thus considered sleep period (S) and wake period (W). Sleep deprivation (SD) was accomplished by presentation of novel objects and gentle cage tapping for 6h starting at light phase onset (from ZT0 to ZT6). Sleep-deprived animals were not allowed to sleep before euthanasia at ZT6. At the indicated timepoints, mice (5 per group for each genotype) were sacrificed by cervical dislocation, their brains rapidly dissected and rinsed in ice-cold PBS, the frontal cortex collected into an icecold tube and flash frozen in liquid nitrogen. Samples were stored at −80°C until further processing (all samples were prepared at the same time).

Frozen cortex tissues were lysed in freshly-prepared 200 μl of urea lysis buffer [8 M urea, 50 Mm NH_4_HCO_3_ supplemented with protease (Roche, 05056489001) and phosphatase inhibitor cocktail (Roche, 04906837001)] at room temperature by mechanical dissociation with pipette. Cortex lysates were sonicated on ice to avoid heating and centrifuged at 17000g for 10min at 4°C. The supernatant was collected and protein concentration was determined by the Pierce BCA protein assay kit (ThermoFisher, 23225). 250μg of each cell lysate were reduced with 5 mM DTT for 1 h at 37 °C and alkylated with 10 mM iodoacetamide for 30 min at room temperature in the dark. The samples were then diluted in 200mM ammonium bicarbonate to reach a final concentration of 1 M urea, and digested overnight at 37 °C with Trypsin/Lys-C (Promega, V5071) at a ratio of 1/50. Digested peptide lysates were acidified with formic acid to a final concentration of 5% formic acid. Samples were centrifuged at 4000 rpm and loaded onto homemade SepPak C18 Tips packed by stacking tree AttractSPE disk (Affinisep, SPE-Disks-Bio-C18-100.47.20) and 2mg beads (Cartridge Waters, 186004621 SepPak C18) into a 200μL micropipette tip for desalting. Peptides were eluted and 90% of the starting material was enriched using Titansphere^™^ Phos-TiO kit centrifuge columns (GL Sciences, 5010-21312) as described by the manufacturer. After elution from the Spin tips, the phospho-peptides and the remaining 10% eluted peptides were vacuum concentrated to dryness and reconstituted in 0.1% formic acid prior to LC-MS/MS phosphoproteome and proteome analyses.

### LC-MS/MS Analysis

Online chromatography was performed with an RSLCnano system (Ultimate 3000, Thermo Scientific) coupled to an Orbitrap Exploris 480 mass spectrometer (Thermo Scientific). Peptides were trapped on a C18 column (75 μm inner diameter × 2 cm; nanoViper Acclaim PepMap™ 100, Thermo Scientific) with buffer A (2/98 MeCN/H2O in 0.1% formic acid) at a flow rate of 3.0 μL/min over 4 min. Separation was performed on a 50 cm x 75 μm C18 column (nanoViper Acclaim PepMap™ RSLC, 2 μm, 100Å, Thermo Scientific) regulated to a temperature of 40°C with a linear gradient of 3% to 29% buffer B (100% MeCN in 0.1% formic acid) at a flow rate of 300 nL/min over 91 min for the phosphoproteome analyses and a linear gradient of 3% to 32% buffer B over 211min for the proteome analyses. MS full scans were performed in the ultrahigh-field Orbitrap mass analyzer in ranges m/z 375–1500 with a resolution of 120 000 (at m/z 200). The top 20 most intense ions were subjected to Orbitrap for phosphoproteomes and top 30 for proteomes for further fragmentation via high energy collision dissociation (HCD) activation and a resolution of 15 000 with the AGC target set to 100%. We selected ions with charge state from 2+ to 6+ for screening. Normalized collision energy (NCE) was set at 30 and the dynamic exclusion to 40s.

### Proteomic and phosphoproteomic analysis

For identification, the data were searched against the Mus Musculus UP000000589 database (downloaded 03/2020) using Sequest HT through Proteome Discoverer (version 2.4). Enzyme specificity was set to trypsin and a maximum of two missed cleavage sites were allowed. Oxidized methionine, N-terminal acetylation, methionine loss and acetylated-methionine loss were set as variable modifications. Phospho serine, threonine and tyrosines were also set as variable modifications in phosphoproteome analyses. Maximum allowed mass deviation was set to 10 ppm for monoisotopic precursor ions and 0.02 Da for MS/MS peaks. False-discovery rate (FDR) was calculated using Percolator (*91*) and was set to 1% at the peptide level for the whole study. The resulting files were further processed using myProMS v.3.9.3 (*92*) (https://github.com/bioinfo-pf-curie/myproms). Label-free quantification was performed using peptide extracted ion chromatograms (XICs), computed with MassChroQ v.2.2.1 (*93*). For protein quantification, XICs from proteotypic peptides shared between compared conditions (TopN matching for proteome setting and simple ratios for phosphoproteome) were used, including peptides with missed cleavages. Median and scale normalization at peptide level was applied on the total signal to correct the XICs for each biological replicate (N=5). The phosphosite localization accuracy was estimated by using the PtmRS node in PD, in PhosphoRS mode only. Phosphosites with a localization site probability greater than 75% were quantified at the peptide level. To estimate the significance of the change in protein abundance, a linear model (adjusted on peptides and biological replicates) was performed, and p-values were adjusted using the Benjamini–Hochberg FDR procedure.

For proteome analyses, proteins were considered in the analysis only when they were found with at least 3 total peptides across 3 biological replicates per group. Then, proteins with an adjusted p-value ≤ 0.05 were considered significantly enriched in sample comparisons. For phosphoproteomic analyses, the phosphoproteome of all analyzed groups was corrected to changes in the proteome before analysis. The peptides threshold was decreased to 1 peptide across 3 biological replicates of a group to consider a phosphosite in the downstream analysis. Then, phosphosites (in one or more phosphopeptides) with an adjusted p-value ≤ 0.05 were considered significantly changed in sample comparisons. Unique phosphosites were also included when identified in at least 3 biological replicates in only one of the groups in each comparison. For analysis of the amplitude of change, the log2 value of the fold change of unique phosphosites was imputed to 5 and −5 (according to the group in which the phosphosite was unique). The density of changed phosphosites per phosphoprotein was determined as the number of significantly changed phosphosites and unique phosphosites identified in one comparison normalized to protein weight (kDa). Proteins and phosphoproteins identified with these criteria were further subjected to Reactome pathway analysis (FDR ≤ 10%) within myProMS (*41*, *42*). The mass spectrometry proteomics and phosphoproteomics data have been deposited to the ProteomeXchange Consortium (http://proteomecentral.proteomexchange.org) via the PRIDE partner repository (*94*) with the dataset identifier PXD030568 (Username; Password: G14n0f2T) For the analysis of proteomic and phosphoproteomic changes on kinases and phosphatases, a list of mouse kinases and phosphatases was generated from Uniprot database by filtering with the following criteria (“kinase” or “phosphatase” and organism “Mus musculus” and “Reviewed”). The lists of kinases and phosphatases obtained were imported into the platform myProMS using their UniProt accession number. Kinases and phosphatases were manually annotated to the different kinase and phosphatases groups using kinhub.org and http://phanstiel-lab.med.unc.edu/coralp/, respectively. Only kinases and phosphatases annotated to the major groups were considered for the analysis. Prediction of kinase activity was done by the web-based tool robust kinase activity inference (RoKAI) at http://rokai.io (*39*). From the phosphoproteomic data comparing micTNF-KO and controls at sleep or wake period, a list of all altered phosphosites, the corresponding protein and the log2 fold change value was created and used as input to RoKAI. Analysis was done using the mouse database as reference, PhosphoSitePlus (PSP) and Signor as kinase substrates databases and the combined KS+PPI+SD+CoEV RoKAI network. From the list of kinases outputs obtained, only kinases with 4 or more substrates and p-value<0.2 were considered.

The lists of SNIPPs and KSPs were imported into myProMS platform using their UniProt accession number and used for analysis of phosphoproteome changes. Synaptic proteins were annotated according to the synaptome database at http://metamoodics.org/SynaptomeDB/index.php (*95*).

### Brain organotypic slices

C57BL/6J pregnant females were obtained from Janvier Labs and P3–P5 pups used for the preparation of organotypic slices as previously described (*96*). Brains were dissected, hemispheres separated in ice-cold dissection medium (33 mM glucose in PBS) and coronal slices (350 μm) cut using a McIllwain tissue chopper (Mickle Laboratory). Slices were plated on Millicell cell culture inserts (Millipore, PICM03050) and maintained in culture medium [MEM (GIBCO, 21090-022) supplemented with 20% heat-inactivated horse serum (GIBCO, 16050-122), 2 mM glutamine, 10 mM glucose, 20 mM HEPES, 10 U/mL penicillin and 10μg/mL streptomycin] at 37°C in 5% CO_2_/air. Culture medium was changed 3 times per week.

Stimulation of organotypic slices with TNFα neutralizing antibody (1 μg/ml, R&D systems, MAB4101) was performed at DIV18-21 in pre-warmed aCSF (aCSF; 125 mM NaCl, 2.5 mM KCl, 2 mM CaCl_2_, 1 mM MgCl_2_, 5 mM HEPES, 33 mM glucose, pH 7.3). For western blot analysis, organotypic slices (3 slices per insert for each condition) were homogenized in cell extraction buffer [10 mM Tris, 100 mM NaCl, 1 mM EDTA, 1% triton-X100, 10% glycerol, 0.1% SDS supplemented with protease (Roche, 05056489001) and phosphatase inhibitor cocktail (Roche, 04906837001)] by mechanical dissociation, centrifuged for 10 min at 16000g at 4°C and the supernatant collected. Total protein concentration was determined by the Pierce BCA protein assay kit (ThermoFisher, 23225).

### Western blot

Equal amounts of protein samples from cortex lysates and organotypic slices lysates were used for immunoblotting. Samples were denatured at 95°C for 5 min in denaturating buffer (125 mM Tris-HCl, 10% glycerol, 2% SDS, bromophenol blue, and 5%β-mercaptoethanol added fresh), resolved by SDS-PAGE in 4-15% Tris-glycine mini-PROTEAN gels (BioRad, 4561086) or homemade 6% polyacrylamide gel (for staining with anti-phospho MARK activation loop antibody) and transferred to PVDF membranes. Membranes were blocked in 5% dry milk, incubated with primary antibody at 4°C overnight and with HRP-conjugated secondary antibody at RT for 1h. Phosphorylated proteins were visualized by chemiluminescence using Lumi-Light Western Blotting substrate (Roche, 12015196001) or SuperSignal West Femto Maximum Sensitivity substrate (Thermo Scientific, 34095). Membranes were scanned on a ImageQuant LAS 4000 imaging system (GE Heathcare). Membranes were stripped with 0.2 M NaOH for 40min and reprobed for loading control. Quantification was performed on Image J. Primary antibodies used were as follows: anti-phospho-MAPK/CDK substrate motif (1:1000, Cell Signaling, 2325), anti-phospho-PKC substrate motif (1:1000, Cell Signaling, 6967), anti-phospho-PKA substrate motif (1:1000, Cell Signaling, 9624), anti-phospho-AKT substrate motif (1:1000, Cell Signaling, 9614), anti-phospho MARK family activation loop (1:1000, Cell Signaling, 4836), anti-vinculin (1:5000, Cell Signaling, 13901) and anti-tubulin (1:10000, Millipore, 05-829).

### Electrode implantation

At 2-4 months of age, under ketamine/xylazine anesthesia, mice were fixed in a stereotaxic apparatus and implanted with electrodes (made of enameled nichrome wire; diameter, 150 μm) for polygraphic sleep monitoring (*97*). Briefly, two electroencephalogram (EEG) electrodes were positioned onto the dura through holes made into the skull over the left frontal cortex and the cerebellum (2 mm lateral to midline and 2 mm anterior to bregma; and at midline, 2 mm posterior to lambda, respectively) and two electromyogram (EMG) electrodes were inserted into the neck muscles. All electrodes were anchored to the skull with Superbond (GACD) and acrylic cement and were soldered to a miniconnector also embedded in cement. Mice were transferred to individual recording cages (20 × 20 × 30 cm) and allowed to recover for 10 days under standard conditions. They were habituated to the recording cables for at least 4 days before recordings were started.

### Sleep recording and sleep deprivation protocol

EMG and EEG signals were recorded with Somnologica software (Medcare, Reykjavik, Iceland), amplified, analog-to-digital converted (2 kHz) and down-sampled at 100 Hz (EMG) or 200 Hz (EEG), and digitalized by an AddLife A/D Module. Undisturbed spontaneous sleep-wake patterns were first examined by recording mice for 24 hr starting at dark onset (baseline recordings). Then, a 6-hr sleep deprivation protocol starting at light onset (ZT0) was achieved by removing the nest, gently moving the mouse, adding new bedding material or novel objects as soon as EEG signs of sleep were detected by the experimenter. The last hour of the protocol (ZT5-6), mice were placed in a new cage to promote exploratory behavior. At ZT6, mice were put back in their home cage and allowed to recover. Matched control recordings (Sham condition) were obtained the day before by gently waking-up mice at ZT6.

### Sleep analysis

Polygraphic recordings were visually scored every 10-s epoch as wake (W), NREM sleep, or REM sleep using the Somnologica software (Medcare, Reykjavik, Iceland). Briefly, wake was defined by low-amplitude/high-frequency EEG signal and high EMG activity; NREM sleep by high-amplitude/low-frequency (<4Hz) EEG signal and reduced EMG activity and REM sleep by dominant theta oscillations (5-10 Hz) on EEG signal and a flat EMG activity with occasional muscle twitches.

The amounts of time spent in each vigilance state during recovery after sleep deprivation or sham condition were expressed as minutes per 3-hr intervals. The sleep architecture was assessed by calculating the mean duration and frequency of vigilance states bouts (a bout could be as short as one epoch).

The EEG signal was processed for power spectrum analysis. Consecutive 10-s epochs were subjected to a fast Fourier transformation routine (FFT), yielding power spectra between 0.4 to 50 Hz, with a 0.4 Hz frequency and a 10-s time resolution. One control mouse for baseline sleep and three control mice for the sleep deprivation experiment were excluded from the spectral analysis as they showed EEG recording artifacts affecting more than 20% of the recording time. For each animal, a mean SWA spectrum corresponding to the 0.5-4.5 Hz frequency band was then obtained by averaging the SWA spectra of an equal number of 10-s epochs of NREM sleep, referred to as quantiles as described in (*98*). The recording sessions were subdivided into 2 quantiles for the baseline dark period, 4 quantiles for the baseline light period and 5 quantiles for the recovery light period immediately following sleep deprivation. The corresponding mean SWA was normalized to the mean SWA during ZT8 to ZT12 of the baseline recording day (*98*).

### Statistical analysis

For western blot data, statistical significance was assessed by non-parametric Mann Whitney test using Prism 8.0 (GraphPad software).

Statistical analyses for sleep data were performed using Prism 9.3 (GraphPad Software). Normality was verified prior to the use of any parametric tests (D’Agostino-Pearson normality test). Data violating normality were log transform. Vigilance states amounts, number and duration of bouts, as well as time-course dynamics of SWA were assessed using two-way repeated-measures analysis of variance (RM-ANOVA). When appropriate, RM-ANOVAs were followed by the Sidak’s multiple comparisons. Statistical significance was considered as P < 0.05, and all results are given as mean values ± SEM.

## Acknowledgments

We gratefully acknowledge Clément Léna for great help with the SWA analysis. We are grateful to Amandine Delecourt, Eleonore Touzalin, Deborah Souchet and the Ibens animal core facility as well as the IBPS Behavioral Core Facility for excellent technical assistance. We thank Patrick Poullet from the bioinformatics platform of the Institut Curie U900 for the continuous development of myProMS and Stéphane Liva for kinase activity analysis made.

## Funding

Agence nationale pour la recherche: SYNTRACK -R17096DJ (AT);

European Research Council: PLASTINHIB Project No: 322821 and MICROCOPS (AT);

Human Brain Project: HBP SGA2-OPE-2018-0017 (AT);

EMBO fellowship: ALTF 362-2017 (MJP);

Région Ile-de-France and Fondation pour la Recherche Médicale (DL);

Marie Skłodowska-Curie Action Individual Fellowship (101031398) from European Commission and National funds through the Portuguese Science and Technology Foundation FCT: PTDC/BIA-CEL/2286/2020 (LFR).

## Author contributions

Conceptualization: AB, AT, MJP, VF.

Formal Analysis: MJP, AB, VF.

Funding acquisition: AB, AT, VF.

Investigation: FJH, LFR, MJP, LC, VF, AB

Software: MJP, FD, VL, DL

Writing – original draft: AB, MJP.

Writing – review & editing: AB, AT, MJP, VF, DL.

## Data and materials availability

All data are available in the main text or the supplementary materials.

