## Supplementary figures for "Microglial TNFα orchestrates brain phosphorylation during the sleep period and controls homeostatic sleep"

**This PDF file includes:**

Figs. S1 to S7
Tables S1 and S5

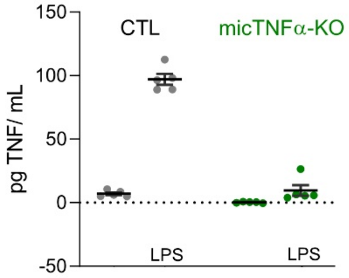


**Figure S1 – Conditional microglia-specific TNFα deletion.**
LPS-induced TNFα release by adult primary microglia observed in control mice (CTL, CX3CR1$^{GFP/+}$:TNF$^{f/f}$) but not on microglia-TNFα depleted mice (micTNFα-KO, CX3CR1$^{CreERT2/+}$:TNF$^{f/f}$). Graph shows TNFα concentration (pg/mL) as mean ± SEM.

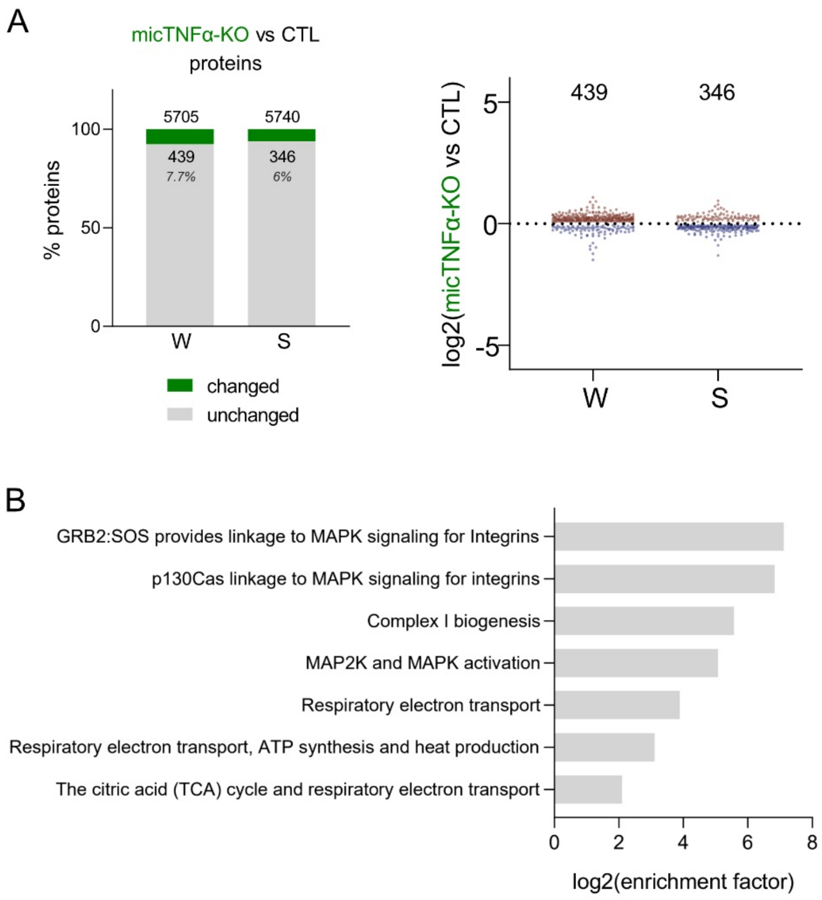


**Figure S2 – Proteome modulation by microglial TNFα during sleep and wake periods.**
(**A**) *Left*, Percentage of proteins showing significant changes in abundance in micTNFα-KO vs CTL comparison during W and S. *Right*, Fold change of significantly different proteins. Each datapoint represents one protein. Datapoints above and below the dotted line show significant up- and downregulation in micTNFα-KO, respectively.
(**B**) Pathways enriched in microglial TNFα-modulated proteome during the sleep period.

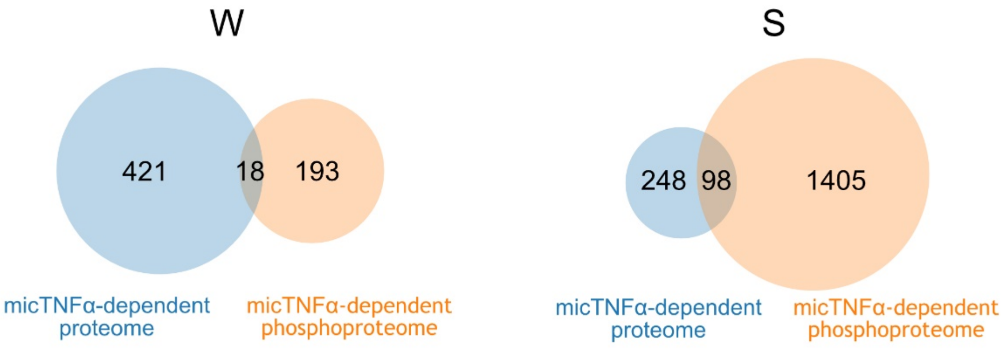


**Figure S3 – Overlap of microglial TNFα-dependent proteome and phosphoproteome during wake and sleep periods.**

Venn diagrams show minimal overlap of significantly changed proteins (blue) and phosphoproteins with at least one changing phosphosite (orange) in micTNFα-KO vs CTL comparison during W and S.

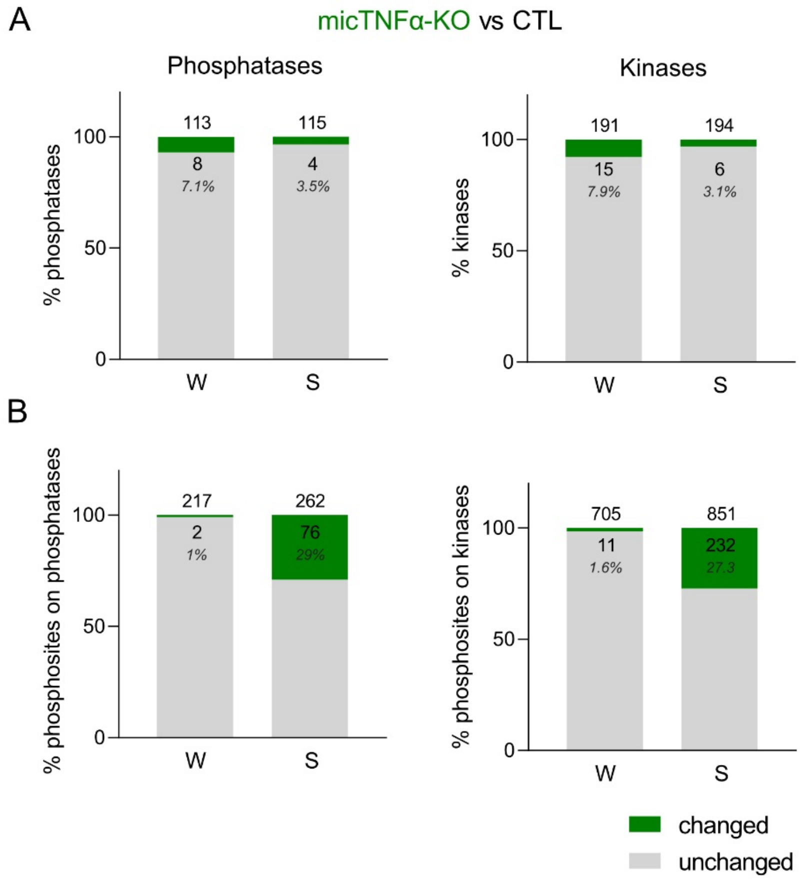


**Figure S4 – Microglial TNFα-dependent proteomic and phosphoproteomic changes in kinases and phosphatases.**

(**A**) Percentage of kinases and phosphatases showing significant changes in abundance in micTNFα-KO vs CTL proteomic comparison during W and S.

(**B**) Percentage of phosphosites on kinases and phosphatases showing significant changes in micTNFα-KO vs CTL phosphoproteomic comparison at W and S.

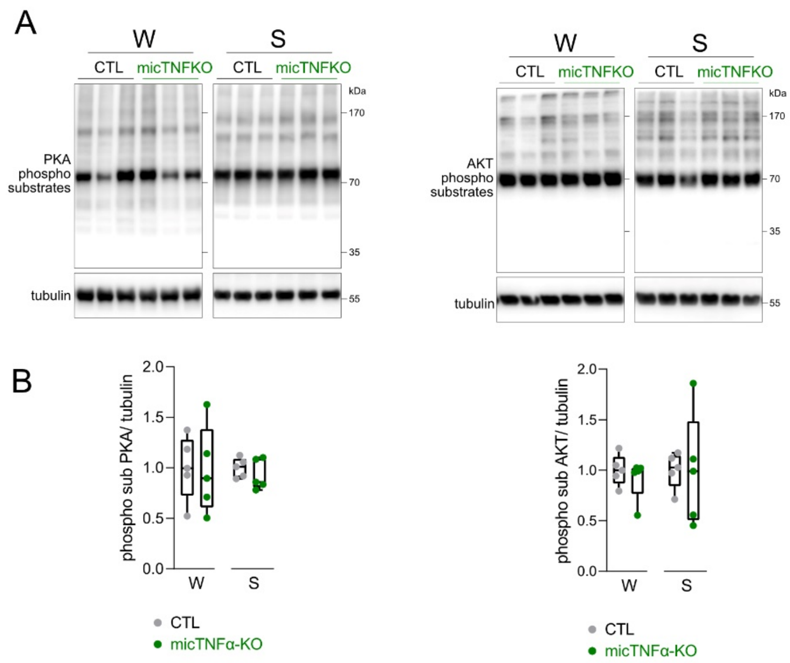


**Figure S5 – Unaltered phosphorylation of PKA and AKT substrates upon loss of microglial TNFα.**

(**A**) Immunoblots of cortical lysates using antibodies specific to PKA and AKT target phosphorylation motifs.

(**B**) Ratio between phosphorylated substrates and loading control normalized to CTL during W and S. n=5 mice per group.

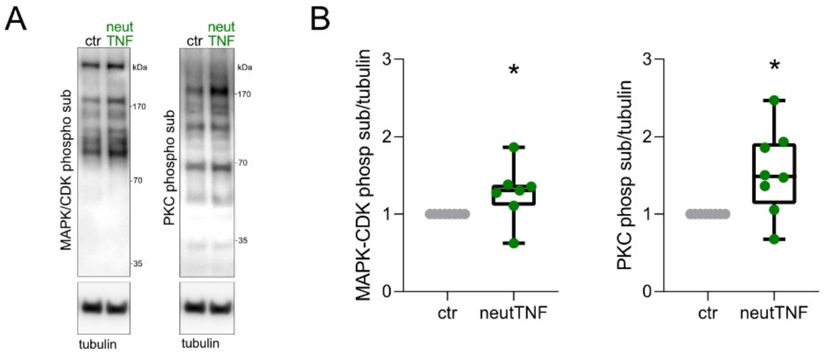


**Figure S6 – Short-term modulation of MAPK/CDK and PKC signaling by neutralization of TNFα on organotypic slices.**

(**A**) Immunoblots of lysates of organotypic slices using antibodies specific to MAPK/CDK and PKC target phosphorylation motifs show effect of neutralizing TNFα antibody (neutTNF). Signal of phosphorylated MARK activation loop on lysates of organotypic slices was too low to be reliably quantified.

(**B**) Ratio between phosphorylated signal of interest and loading control normalized to control. n= 7-8 independent experiments. *$p<0.05$, Mann Whitney test.

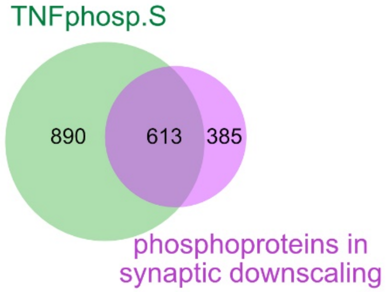


**Figure S7 – Phosphosubstrates of synaptic downscaling are targets of microglial TNFα during the sleep period.**

Venn diagram shows high overlap of TNFα-modulated phosphoproteins during the sleep period (TNFphosp.S) and proteins undergoing phosphorylation changes upon bicuculine-induced synaptic downscaling (at 5min, 15min and 24h) (75).

**Table S1. (separate file)**
Daily oscillations in cortical phosphoproteome

**Table S2. (separate file)**
Microglial TNFα-dependent changes in phopshoproteome at W and S

**Table S3. (separate file)**
Microglial TNFα-dependent changes in proteome at W and S

| | | Number of bouts | | Bouts mean duration (min) | |
|---|---|---|---|---|---|
| | | **Sham** | **SD** | **Sham** | **SD** |
| **WAKE** | CTL | 42.2 ± 2.2 | 39.8 ± 1.9 | 1.35 ± 0.14 | 0.97 ± 0.06 |
| | micTNFα-KO | 45.2 ± 2.6 | 41.3 ± 2.6 | 1.15 ± 0.14 | 1.20 ± 0.10 |
| **NREM sleep** | CTL | 42.2 ± 2.2 | 40.1 ± 1.9 | 2.71 ± 0.98 | 3.27 ± 0.16** |
| | micTNFα-KO | 45.4 ± 2.6 | 41.6 ± 2.6 | 2.59 ± 0.13 | 3.01 ± 0.19 |
| **REM sleep** | CTL | 13.4 ± 1.2 | 13.1 ± 0.6 | 1.17 ± 0.07 | 1.21 ± 0.05 |
| | micTNFα-KO | 17.3 ± 1.0$^{##}$ | 11.9 ± 0.8*** | 1.11 ± 0.07 | 1.20 ± 0.09 |

**Table S4. Sleep and wake characteristics in CTL and micTNFα-KO mice in the first 3 hr of the recovery phase after sleep deprivation (SD) or Sham condition (Sham).**
The data are given as means ± S.E.M (n=15). For multiple group comparisons: $^{**}$P < 0.01 and $^{***}$P < 0.001 Sham versus SD and $^{##}$P<0.01 CTL versus micTNFα-KO. For statistics see table S5.

| Figure # Table # | Panel | | n | Test used | Factor | F/t value | P value |
|---|---|---|---|---|---|---|---|
| **Figure 5** | **A** | SWA in baseline sleep | 14/15 | 2-way RM ANOVA | Genotype<br>Time<br>GxT | F(1,27) = 3.761<br>F(1.813,75.95) = 181.0<br>F(5,135) = 3.108 | P=0.0630<br>P<0,0001<br>P=0.0109 |
| | | SWA after SD | 12/15 | 2-way RM ANOVA | Genotype<br>Time<br>GxT | F(1,25) = 1.257<br>F(3.271,81.78) = 281.3<br>F(4,100) = 4.044 | P=0.2729<br>P<0,0001<br>P=0.0044 |
| | **B** | Wake | 15 | 2-way RM ANOVA | Genotype<br>SD<br>GxSD | F(1,28) = 0.4188<br>F(1,28) = 6.435<br>F(1,28) = 4.686 | P=0.5228<br>P=0,0170<br>P=0.0391 |
| | | NREMS | 15 | 2-way RM ANOVA | Genotype<br>SD<br>GxSD | F(1,28) = 0.8128<br>F(1,28) = 10.965<br>F(1,28) = 2.518 | P=0.3750<br>P=0,0026<br>P=0.1238 |
| | | REMS | 15 | 2-way RM ANOVA | Genotype<br>SD<br>GxSD | F(1,28) = 0.4485<br>F(1,28) = 6.046<br>F(1,28) = 11.99 | P=0.5085<br>P=0,0204<br>P=0.0017 |
| **Table S4** | **Number** | Wake | 15 | 2-way RM ANOVA | Genotype<br>SD<br>GxSD | F(1,28) = 0.7771<br>F(1,28) = 2.304<br>F(1,28) = 0.1262 | P=0.3855<br>P=0.1402<br>P=0.7251 |
| | | NREMS | 15 | 2-way RM ANOVA | Genotype<br>SD<br>GxSD | F(1,28) = 7403<br>F(1,28) = 2.104<br>F(1,28) = 0.1344 | P=0.3969<br>P=1580<br>P=0.7167 |
| | | REMS | 15 | 2-way RM ANOVA | Genotype<br>SD<br>GxSD | F(1,28) = 1.163<br>F(1,28) = 10.49<br>F(1,28) = 11.84 | P=0.2901<br>P=0031<br>P=0.0018 |
| | **Mean** | Wake | 15 | 2-way RM ANOVA | Genotype<br>SD<br>GxSD | F(1,28) = 0.024<br>F(1,28) = 1.670<br>F(1,28) = 2.719 | P=0.8782<br>P=0.2069<br>P=0.1104 |
| | | NREMS | 15 | 2-way RM ANOVA | Genotype<br>SD<br>GxSD | F(1,28) = 0.9717<br>F(1,28) = 14.56<br>F(1,28) = 0.3350 | P=0.3327<br>P=0.0007<br>P=0.2764 |
| | | REMS | 15 | 2-way RM ANOVA | Genotype<br>SD<br>GxSD | F(1,28) = 0.5964<br>F(1,28) = 1.051<br>F(1,28) = 0.026 | P=0.4464<br>P=0.3141<br>P=0.8737 |

**Table S5. Statistical analysis for figure 5 and table S4.**
Statistical tests for each figure/table and panel, with the test used, the number of mice (n) and the F/t value.
